# Patterns of polymorphism, selection and linkage disequilibrium in the subgenomes of the allopolyploid *Arabidopsis kamchatica*

**DOI:** 10.1101/248195

**Authors:** Timothy Paape, Roman V. Briskine, Heidi E.L Lischer, Gwyneth Halstead-Nussloch, Rie Shimizu-Inatsugi, Masaomi Hatekayama, Kenta Tanaka, Tomoaki Nishiyama, Renat Sabirov, Jun Sese, Kentaro K. Shimizu

**Affiliations:** Department of Evolutionary Biology and Environmental Studies and Department of Plant and Microbial Biology, University of Zurich, Winterthurerstrasse 190, CH-8057 Zurich, Switzerland; Functional Genomics Center Zurich, Winterthurerstrasse 190, CH-8057 Zurich, Switzerland; Swiss Institute of Bioinformatics (SIB), Lausanne, Switzerland; Sugadaira Montane Research Center, University of Tsukuba, Ueda, Nagano, Japan; Advanced Science Research Center, Kanazawa University, 13-1 Takara-machi, Kanazawa, 920-0934, Japan; Institute of Marine Geology and Geophysics, Far East Branch, Russian Academy of Sciences, Yuzhno-Sakhalinsk, Russia; Artificial Intelligence Research Center, National Institute of Advanced Industrial Science and Technology (AIST), Tokyo, Japan; Kihara Institute for Biological Research, Yokohama City University, 641-12 Maioka, 244-0813 Totsuka-ward, Yokohama, Japan.

## Abstract

Although genome duplication is widespread in wild and crop plants, little is known about genome-wide selection due to the complexity of polyploid genomes. In allopolyploid species, the patterns of purifying selection and adaptive substitutions would be affected by masking owing to duplicated genes or ‘homeologs’ as well as by effective population size. We resequenced 25 distribution-wide accessions of the allotetraploid *Arabidopsis kamchatica*, which has a relatively small genome size (450 Mb) derived from the diploid species *A. halleri* and *A. lyrata*. The level of nucleotide polymorphism and linkage disequilibrium decay were comparable to *A. thaliana*, indicating the feasibility of association studies. A reduction in purifying selection compared with parental species was observed. Interestingly, the proportion of adaptive substitutions (α) was significantly positive in contrast to the majority of plant species. A recurrent pattern observed in both frequency and divergence-based neutrality tests is that the genome-wide distributions of both subgenomes were similar, but the correlation between homeologous pairs was low. This may increase the opportunity of different evolutionary trajectories such as in the *HMA4* gene involved in heavy metal hyperaccumulation.

## Introduction

Genome duplication is a widespread evolutionary force in plants. As many as 35% of vascular plants are recent polyploid species^1^ and increased ploidy is particularly common in crops^2^. The abundance of polyploid species in plants motivated speculation and theoretical analysis on the advantages and disadvantages of genome duplication^3,4^. However, compared with diploid species, much less is known about the genome-wide patterns of polymorphism and selection due to the complexity of polyploid genomes^5^. Major difficulties in genome scale analyses result from the large genome sizes of polyploids and the high similarity between the duplicated chromosomes. However, recent advances in next-generation sequencing and bioinformatic tools^6,7^ are enabling genome-wide data to study polymorphisms and transcriptomics patterns for entire subgenomes in newly emerging model polyploids^8–11^.

Genome-wide strengths of positive and purifying selection can be quantified using several complementary approaches. Frequency-based tests using site-frequency spectra (SFS) such as Tajima’s *D* and Fay and Wu’s *H* statistics can detect rare or common polymorphisms that are due to purifying or positive selection. Divergence-based tests compare interspecific divergence (from an outgroup) to intraspecific polymorphism to identify positive selection on amino-acid substitutions^12^. These tests include several derivatives of the McDonald-Kreitman test^13^ or “MK-tests”, such as the direction of selection (DoS) neutrality index^14^, and methods to estimate the distribution of fitness effects (DFE) and proportion of adaptive substitutions (α)^13^ in genome-wide data. Theoretical and empirical studies in plant species using these methods^15,16^ showed that the strengths of selection are affected by species-specific characteristics such as the effective population size (*N_e_*), mating system, and genome duplication, which are mutually interacting. In particular, species with low *N_e_* typically have the highest proportions of neutral mutations^15,17^, while species with large *N_e_* have higher proportions of non-synonymous substitutions under purifying selection and adaptive evolution^8,19,20^.

Allopolyploidization should have a profound effect on the patterns of polymorphism and selection. First, the redundancy of duplicated gene copies of similar function from different parents (“homeologs”) may affect the strength of selection. At the early stages, genome duplication may increase evolutionary rates of duplicated genes^21,22^ and may facilitate the evolution of new adaptive function because the original function can be retained in other copies (so-called neofunctionalization model)^23,24^. In contrast, the additional copy may mask the effect of adaptive and deleterious mutations^4,16^. Second, polyploidization must involve a reduction in *N*_e_ due to a bottleneck during speciation. In addition, polyploid speciation is typically associated with the transition from outcrossing to self-fertilization, which reduces *N_e_* several times less than parental species (at least half)^25^. While studies of selection in polyploids are very limited, a recent empirical study in the allotetraploid *Capsella bursa-pastoris* showed a decrease in the efficacy of purifying selection in one of the subgenomes but an increase in another subgenome^8^. Further empirical studies are necessary to compare the consequences of genome duplication in polyploid species.

The genus *Arabidopsis* has both auto- and allopolyploid species in addition to the more well-studied diploid relatives^26^. *Arabidopsis kamchatica*^27^ is a recent allopolyploid (estimated 20,000-250,000 years ago)^28^ derived from the two diploid species *A. halleri* (particularly subsp. *gemmifera* distributed in East Asia), and *A. lyrata* (particularly subsp. *petraea* from Far East Russia)^29–31^. The two diploid parents are predominantly self-incompatible (SI) while a transition to selfing accompanied allopolyploid formation^28^. The genome size (about 450 Mb) is relatively small among polyploid species^32,33^ which is an advantage for resequencing. The species distribution of *A. kamchatica* is very broad, ranging from Taiwan, Japan, Far East Russia, Alaska and Pacific Northwest, USA. The high variation in latitude and altitude compared with the parental species^34,35^ suggests that merging the diploid transcriptional networks and parental adaptations provided the allopolyploid with plasticity to inhabit diverse environments^10^.

To understand the ecological distributions of polyploids, genetically tractable traits are essential. Heavy metal tolerance and hyperaccumulation likely influenced ecological divergence and speciation between the parental species of *A. kamchatica* (*A. halleri* and *A. lyrata*) due to adaptive mutations in metal transporter genes such as *HEAVY METAL ATPASE4* (*HMA4*)^36,37^. The *HMA4* locus has been shown to be the primary transporter of cadmium and zinc from roots to shoots in *A. halleri* due to a tandem triplication and enhanced *cis*-regulation, while only a single copy of *HMA4* exists in the non-hyperaccumulators *A. lyrata* and *A. thaliana*. *A. kamchatica* inherited hyperaccumulation from the diploid parent *A. halleri*, although attenuated expression of *halleri-*derived *HMA4* and putatively inhibiting *lyrata*-derived factors reduced the trait to about half of *A. halleri*^10^. Estimates of genetic diversity surrounding the *HMA4* region in *A. halleri* suggests a hard selective sweep^38^ which may have predated the formation *A. kamchatica*^10^.

Here, we used *de novo* assemblies of the closest diploid relatives of *A. kamchatica* to sort Illumina reads to their respective subgenomes using a distribution-wide collection of 25 natural allopolyploid accessions. We used population genomics to ask: a) what is the level of genome-wide diversity compared with diploid outcrossing and selfing *Arabidopsis* species? b) are there differences in polymorphism, allele frequencies, linkage disequilibrium (LD), and selection between subgenomes? c) do pairs of homeologs tend to show similar patterns in diversity and neutrality? d) does the *HMA4* locus show significant differences in genetic diversity between homeologs and how does diversity surrounding this locus compare with the genome-wide average? e) what proportions of the subgenomes show neutral, deleterious, or adaptive mutations and how do they differ from the diploid parents? and f) are there high frequencies of loss of function mutations in either subgenome? Together, these plant accessions and polymorphism data will serve as a core diversity panel for further studies of genotype-phenotype associations and the genetic architecture of complex traits using larger collections of globally collected samples.

## Results

### Reference Genome Assembly and Allopolyploid Resequencing

To sort Illumina reads of *A. kamchatica* to their parentally-derived subgenomes, we generated long mate-pair *de novo* assemblies of *A. lyrata* subsp. *petraea* (also called *A. petraea* subsp. *umbrosa*) in addition to East Asian *A. halleri* subsp. *gemmifera* which we previously reported^39^. Assembly statistics indicated that the *A. lyrata* and *A. halleri* reference genomes have scaffold N50 of 1.2 Mb and 0.7 Mb, comprising 1,675 and 2,239 scaffolds respectively (Table 1, Supplementary Table 1 and 2 for gene annotation statistics), providing opportunities to compare diversity over very large syntenic regions in the allopolyploid subgenomes.

**Table 1.** Reference genome assembly statistics of v2.2 of Siberian *A. lyrata* subsp. *petraea* and v2.2 of *A. halleri* subsp. *gemmifera* (Tada Mine).

| Assembly statistics | *A. lyrata* | *A. halleri*<sup>a</sup> |
| --- | --- | --- |
| Length (bp) | 175,182,717 | 196,243,198 |
| Missing (%)<sup>b</sup> | 12.75 | 14.81 |
| Scaffolds | 1,675 | 2,239 |
| Shortest scaffolds (bp) | 940 | 932 |
| Longest scaffolds (bp) | 6,771,235 | 4,302,264 |
| Scaffold N50 length (bp) | 1,260,070 | 712,249 |
| Scaffold N50 count | 38 | 71 |
| NG50 length (bp) | 804,357 | 489,153 |
| NG50 count | 63 | 117 |
| Genome size (FC)<sup>c</sup> | 225 Mb | 250 Mb |

a: *A. halleri* assembly previously reported in Briskine et al.<sup>39</sup>
b: Missing data counted as number of N's in the assembly and is percentage of total length
c: Genome size measured by flow cytometry (FC)

We sorted reads of 25 individuals from a distribution-wide collection (Supplementary Table 3) of *A. kamchatica* to their parental origins by first aligning each read to both parental genomes then classified the reads as ‘*origin*’ reads (*halleri*-derived = H-origin, *lyrata*-derived = L-origin) using algorithms that quantify mismatches to either parent^32^. Our accessions had on average 12.5X coverage for the H-origin-subgenome (range 5.2X - 20.7X) and on average 10.7X coverage for the L-origin-subgenome (range 4.3X - 17.7X). Homeolog specific PCR and Sanger sequencing was used to validate SNPs and read sorting for twelve genes and showed that reads were accurately assigned to their respective subgenomes (Supplementary Material). In addition, pyrosequencing was used in two previous studies to detect ratios of parentally derived SNPs to validate homeolog specific expression (RNA-seq) in ten other genes^10,32^ where the same read sorting pipeline was used. After filtering for SNP quality and coverage, our resequencing dataset resulted in 1,674,191 H-origin and 1,930,341 L-origin SNPs. Using the parental genome assemblies for *A. kamchatica* SNP calling, we identified ca. 23,500 homeologous coding sequences using reciprocal best BLAST hits (Supplementary Table 2), of which ca. 21,500 show orthology to *A. thaliana*, representing 72% and 67% of our annotated genes respectively.

### Genome-wide Nucleotide Diversity in *A. kamchatica*

We examined the patterns of nucleotide diversity for ca. 21,000 coding sequences of both *halleri* and *lyrata*-derived homeologs in *A. kamchatica* that could be aligned to *A. thaliana* orthologs as the outgroup. We found that both subgenomes showed similar mean values of nucleotide diversity (π) (π _coding_ = 0.0014 bp^-1^ for *halleri*-subgenome and π_coding_ = 0.0015 bp^-1^ for *lyrata*-subgenome, and π_coding_ = 0.0015 bp^-1^ when combined) although the *lyrata*-derived homeologs showed slightly broader ranges in π (Table 2, Fig. 1A). Nucleotide diversity at synonymous sites (π_*syn*_) was also similar for the two subgenomes with a slightly higher value for the *lyrata-*subgenome (π_*syn*_ = 0.0049) than in the *halleri*-subgenome (π_*syn*_ = 0.0044). The nucleotide diversity in *A. kamchatica* is about six times lower than European *A. halleri* and *A. lyrata* (π_*syn*_ = 0.029 for *A. halleri* and 0.028 for *A. lyrata* estimated using resequencing data from^30^) and is more similar to that of *A. thaliana* (π_*syn*_ = 0.0059 - 0.007)^17,30,40^. Sliding window analysis including non-coding regions also showed comparable values (Supplementary Table 4).

**Fig. 1.**
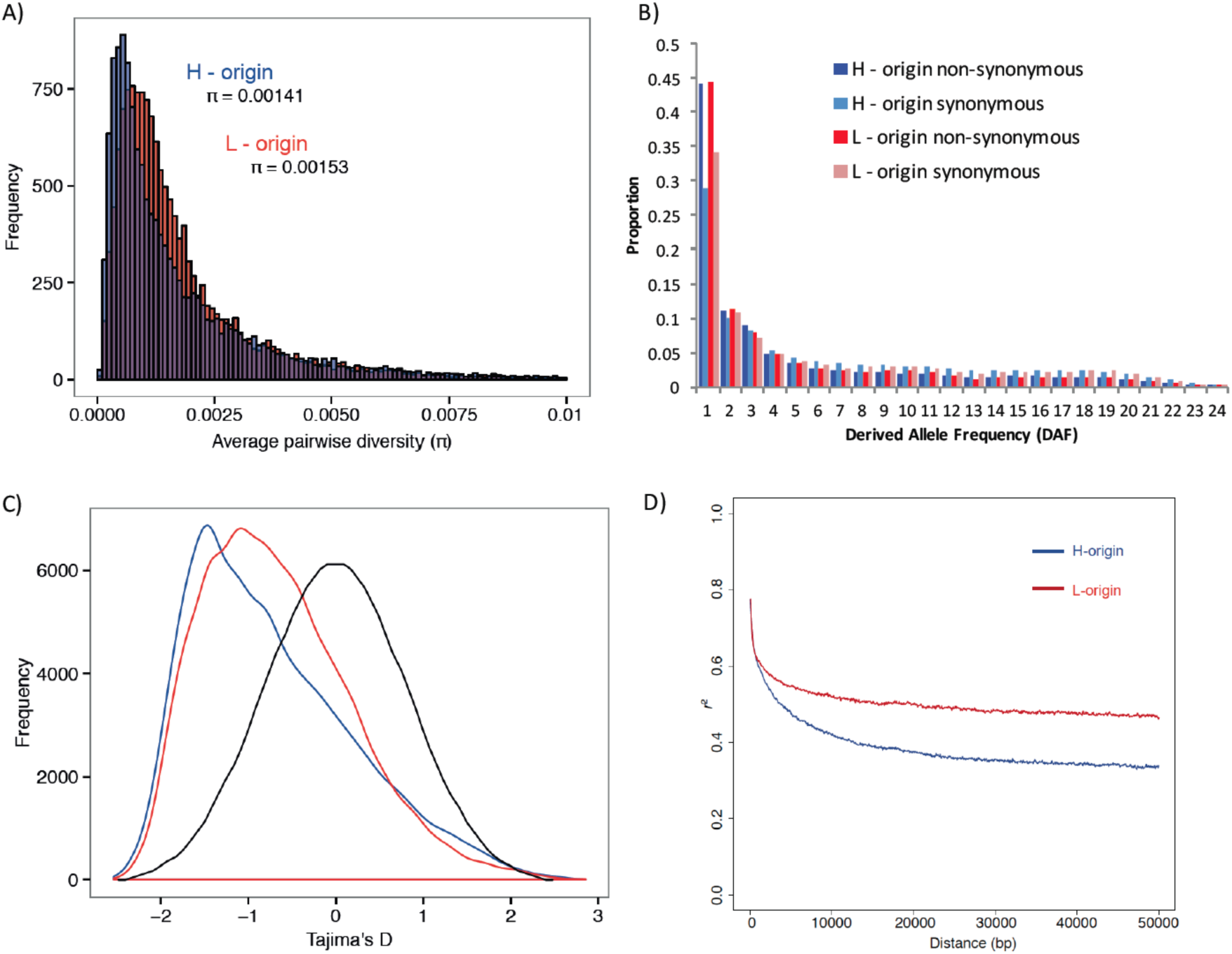
Genome-wide diversity and linkage disequilibrium. (A) Average pairwise diversity (π) of *halleri*-origin (H-origin) and *lyrata-*origin (L-origin) coding sequences. (B) H-origin and L-origingenes show no significant differences in proportions of non-synonymous and synonymous substitutions (*X*^2^, p-value = 0.58), and the majority of substitutions are low at frequency. (C) Tajima’s D distributions for both genomes (blue density curve = H-origin, red density curve = L-origin) show departures from neutrality (black density curve where neutral = 0), mean values for both distributions are negative (Table 1). (D) The mean decay of linkage disequilibrium (LD) estimated using 100 kb sliding windows shows mean LD decay < 10 kb for both H-origin (blue) and L-origin (red) genomes.

**Table 2.**
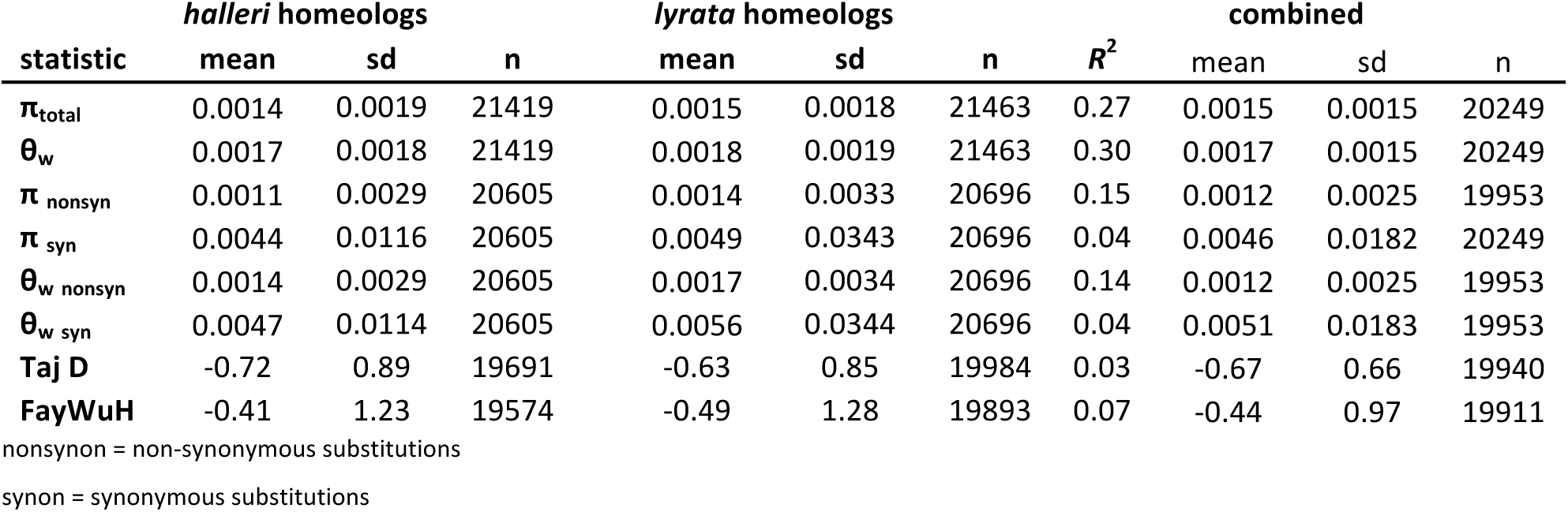
Diversity statistics for coding sequences (CDS) of *A. kamchatica* homeologs. Values are average pairwise diversity, π, polymorphism Watterson’s estimator, θ_w_, Tajima’s D, Fay and Wu’s H. Correlations between homeolog diversity stastistics are shown as *R*^**2**^ correlation coefficient.

We calculated the effective population size, *N*_*e*_, using our empirical estimates of π for *A. kamchatica* and both diploid species and two different mutation rates^41,42^. The estimated values for *A. kamchatica* were: *N*_*e*_ = 77,000 and 54,000 using the two mutation rates respectively. The values for *A. kamchatica* were several times lower than *A. halleri*: *N*_*e*_ = 467,000 and 364,000 and *A. lyrata*: *N*_*e*_ = 483,000 and 345,000 (Supplementary Table 5). We interpret these estimates of *N*_*e*_ with caution as the mutation rates for these species have not been estimated directly and the diversity estimates used in the calculation can themselves be affected by demography. The estimates are nevertheless useful as general comparisons between species to identify large differences in magnitude^17,19^.

Higher proportions of non-synonymous mutations were found to be at low frequency compared with synonymous mutations and no significant differences in the relative proportions were found between subgenomes (Fig. 1B). This suggests purifying selection on a large proportion of amino-acid changing substitutions in both subgenomes. Frequency-based test statistics clearly show significant departures from neutrality for both subgenomes (Fig. 1C). The mean values of Tajima’s D were negative for both subgenomes (Table 2, Fig. 1C) owing to high proportions of rare variants.

The distributions and means of Tajima’s D in *A. kamchatica* (Table 2) are similar to early genome-wide data from *A. thaliana* (mean Tajima’s D_*A.thaliana*_ = -0.8)^43^, although more recent estimates using over 300 genomes show a higher mean but not higher median in *A. thaliana* (mean D_*A.thaliana*_ = 0.006, median D_*A.thaliana*_ = -0.33)^30^, which likely reflects more intermediate-frequency polymorphisms in the large species wide sample. The same study^30^ reported an excess of rare variants in the diploid relatives of *A. kamchatica* (mean D_*A.lyrata*_ to be -0.99 in *A. lyrata* and D_*A.halleri*_ = -0.23 in *A. halleri*).

We found the means of the distributions for most summary statistics to be very similar between the two subgenomes, but when pairs of all homeologs were compared correlations were generally low for diversity and neutrality estimators (Table 2). The correlations of π _syn_ and θ_w syn_ were both nearly zero (Table 2). Similarly, the distributions and means of Tajima’s D overlap for both subgenomes but the correlation for Tajima’s D between pairs of homeologs is very low (*R*^2^ = 0.03). The Fay and Wu’s H statistic, which detects departures from neutrality due to intermediate and high frequency variants, also shows a very low correlation between homeologs (Table 2). Higher correlations were observed for non-synonymous or total sites, but this can be explained by the constraints on non-synonymous changes. In summary, the low correlations are consistent with different evolutionary trajectories of individual homeologous pairs.

### Mean Rate of LD Decay in Both Subgenomes is Similar But Not Equal

Long scaffold assemblies allowed us to estimate genome-wide LD for each subgenome to evaluate the feasibility of association mapping in *A. kamchatica*. We found that mean LD decay was between 5-10 kb for both subgenomes (Fig. 1D), which is similar to the self-fertilizing species *A. thaliana* and *M. truncatula* which show LD decay within 2-10 kb ranges^44,45^. The mean LD for the *lyrata*-subgenome decayed slightly faster and remained at *r*^2^ = 0.47 over the scale of > 100 kb genomic regions while mean LD for the *halleri-*subgenome leveled off at *r*^2^ = 0.34 > 100 kb. The 50% and 90% confidence intervals around the mean LD decay also revealed much greater variance in the *lyrata*-subgenome (Supplementary Fig. 1).

Population structure assignments and phylogenetic clustering may provide some explanation for subgenome differences in LD. The 25 accessions cluster geographically with one main clade/group comprising the northern accessions (Russia, Sakhalin, and Alaska) and the other main group containing Japanese accessions (Supplementary Fig. 2,3). The branch lengths within these groups for the *lyrata*-subgenome are shorter than for the *halleri*-subgenome, particularly in the Japanese clade, indicating greater relatedness. These clusterings are also consistent with previous haplotype analysis using low density nuclear and chloroplast markers^29^.

### Diversity of the *HMA4* Locus and the Genomic Background

We analyzed genetic diversity on the scaffolds containing the *HMA4* locus to compare whether it differs from the genomic background and the surrounding regions flanking the *HMA4* coding sequences. We centered the main genomic region containing the *HMA4* coding sequences which we call “HMA4-M” (containing 17 coding sequences). This region spans 304 kb on *A. halleri* (scaffold_116) and spans 155 kb on *A. lyrata* (scaffold_52). While the differences in length of HMA4-M between the parental genomes can be attributed to the triplicated *HMA4* genes in *A. halleri*, the genes surrounding *HMA4* in both *A. halleri* and *A. lyrata* are syntenic (Fig. 2A). To compare HMA4-M to surrounding regions, we used the upstream adjacent region (left-side) “HMA4-L” (containing 13 coding sequences) which is 125 kb for the *A. halleri* region and 183 kb in *A. lyrata*, and the downstream adjacent region (right-side) “HMA4-R” (containing 13 CDS sequences), which is 105 kb in the *A. halleri* region and ca. 50 kb for *A. lyrata*.

**Fig. 2.**
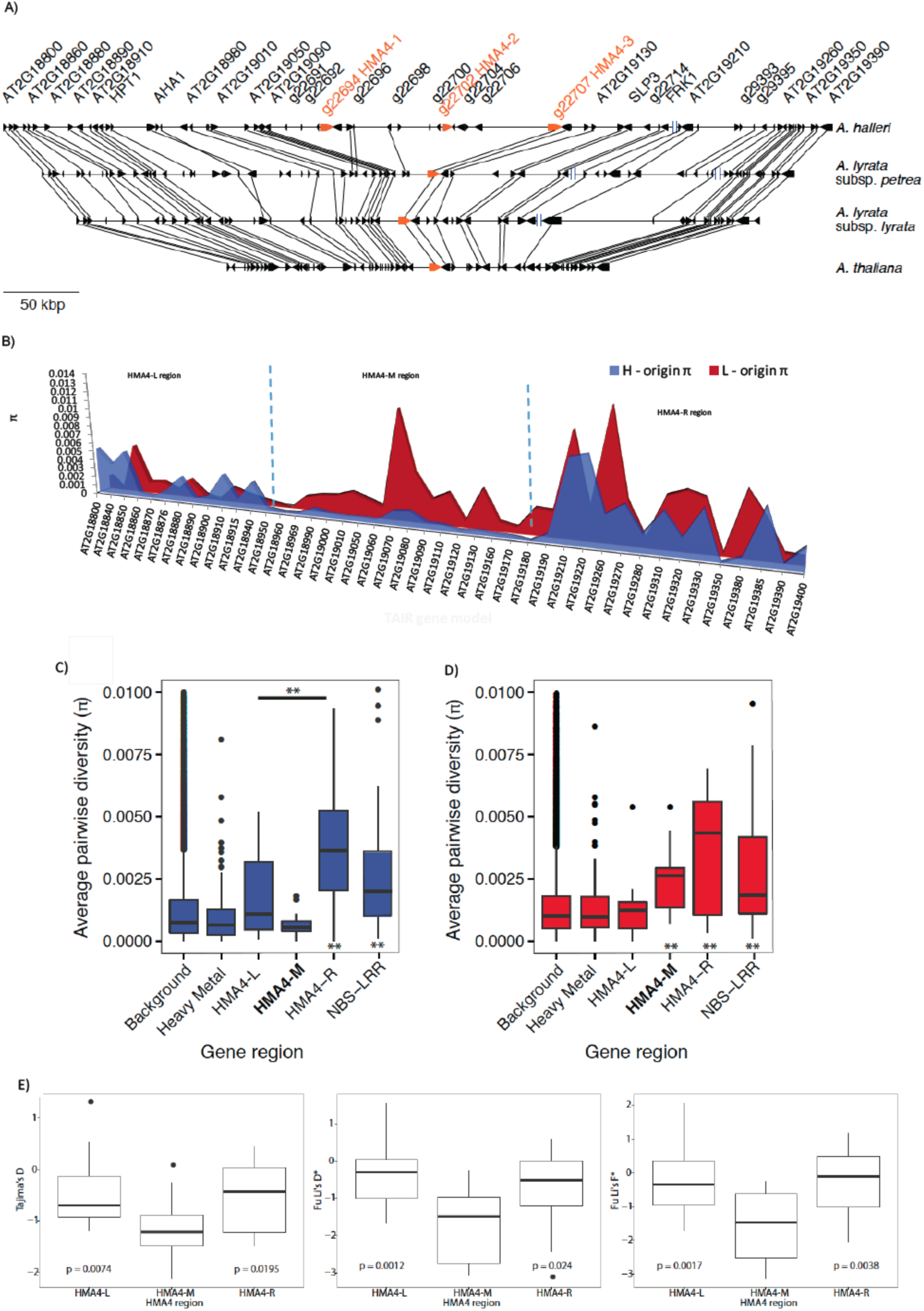
Genetic diversity of the syntenic *HMA4* region. (A) Synteny of the *HMA4* region from *A. halleri* v2.2^39^, *A. lyrata* subsp. *petraea* v2.2, *A. lyrata* subsp. *lyrata* (JGI)^70^ and *A. thaliana* (TAIR). Average pairwise diversity (π) of genes surrounding the *HMA4* region in both homeologs of *A. kamchatica*. (C) For the *halleri*-subgenome, genetic diversity of NBS-LRRs is significantlygreater (two asterisks below, ^**^p < 0.001) than diversity compared with the background while heavy metal (HM) genes show no significant difference. Diversity for both HMA4-L (π = 0.0018) and HMA4-R (π = 0.004) are significantly higher than the HMA4-M (which contains the *HMA4* coding sequences) region (two asterisks above HMA4-M, ^**^p < 0.001). (D) For the *lyrata*-subgenome, diversity of NBS-LRRs, HMA4-M and HMA4-R are all significantly higher than the background. The diversity of the *lyrata* HMA4-M (π = 0.0032) region is also significantly greater than the *halleri* HMA4-M region (π = 0.0007, paired *t-test* p-value = 0.003; Wilcoxon sign rank p-value = 0.0001). The neutrality statistics (E) Tajima’s D, Fu and Li’s D^*^ and Fu and Li’s F^*^ all show the *halleri*-origin HMA4-M region to be significantly lower than the left and right flanking regions supporting genetic hitchhiking surrounding the *HMA4* coding sequences.

The distribution of π in the HMA4-M region for H-origin genes showed low diversity (π_mean_ = 0.0007) but it is not significantly lower than the background genes (Fig. 2B and 2C). However, the two adjacent regions (HMA4-L and HMA4-R) compared to the HMA4-M (containing the *HMA4* coding sequences) region have significantly greater diversity (Fig. 2B and 2C). Furthermore, we found significantly lower Tajima’s D, Fu & Li’s D^*^ and Fu & Li’s F^*^ statistics in the HMA4-M region compared with both adjacent regions (Fig. 2E), suggesting greater selection on the HMA4-M region. The significantly lower diversity and neutrality statistics in HMA4-M compared with the adjacent regions likely defines the window of the sweep region previously reported for *A. halleri* ^38^.

Unlike the *halleri* HMA4-M region, the diversity of the *lyrata* HMA4-M region is significantly greater than the genomic background (p-value = 0.0028), but not different from the two adjacent regions (Fig. 2D). Moreover, the *lyrata*-HMA4-M region shows no significant differences from the adjacent HMA4-L or HMA4-R regions for Tajima’s D, Fu & Li’s D^*^ and Fu & Li’s F^*^ (not shown). The elevated diversity of the *lyrata*-origin *HMA4* locus compared with the genomic background is consistent with relaxed selective constraint on the *lyrata*-origin *HMA4* locus.

We also estimated diversity of all annotated heavy metal transporters, metal ion transporters, and metal homeostasis genes for comparison with the genome-wide average (HM genes, N=118 genes). We expected these genes to have low overall diversity in both genomes due to selective constraint as many of these ion transporters are expected to have roles in basic metal homeostasis^46^. As a contrast, we compared NBS-LRR genes (N=39 genes) which have putative roles in plant defense and have high diversity in plants^47,48^ and are expected to have equally high diversity in both subgenomes. The HMA4-L and HMA4-R regions in both subgenomes have more similar levels of diversity to NBS-LRR’s than to the genomic background or HM genes (Fig. 2C and 2D).

### The Majority of Homeologous Proteins Showed Signatures of Purifying Selection

Next we employed divergence-based tests to estimate the strength of purifying and positive selection on amino-acid changing substitutions. We calculated the divergence of each homeolog from the outgroup *A. thaliana* to estimate the relative proportions of diverged non-synonymous (*D*_n_) and synonymous (*D*_s_) sites to polymorphic non-synonymous (*P*_n_) and synonymous (*P*_s_) sites. For each gene, The counts of *D*_n_, *D*_s_, *P*_n_, and *P*_s_ for the coding regions of both subgenomes were used to estimate the direction of selection (DoS)^14^, a neutrality index that varies from -1.0 to 1.0, where zero indicates neutrality and negative and positive values indicate purifying and positive selection, respectively. Both subgenomes had similar distributions in DoS with means of -0.2 (Fig. 3A) suggesting that 68-71% of proteins derived from both subgenomes are under purifying selection (when DoS is < -0.01). Like the previous summary statistics, the correlation in DoS between *halleri* and *lyrata* homologs is positive but fairly low (*R*^2^ = 0.17).

**Fig. 3.**
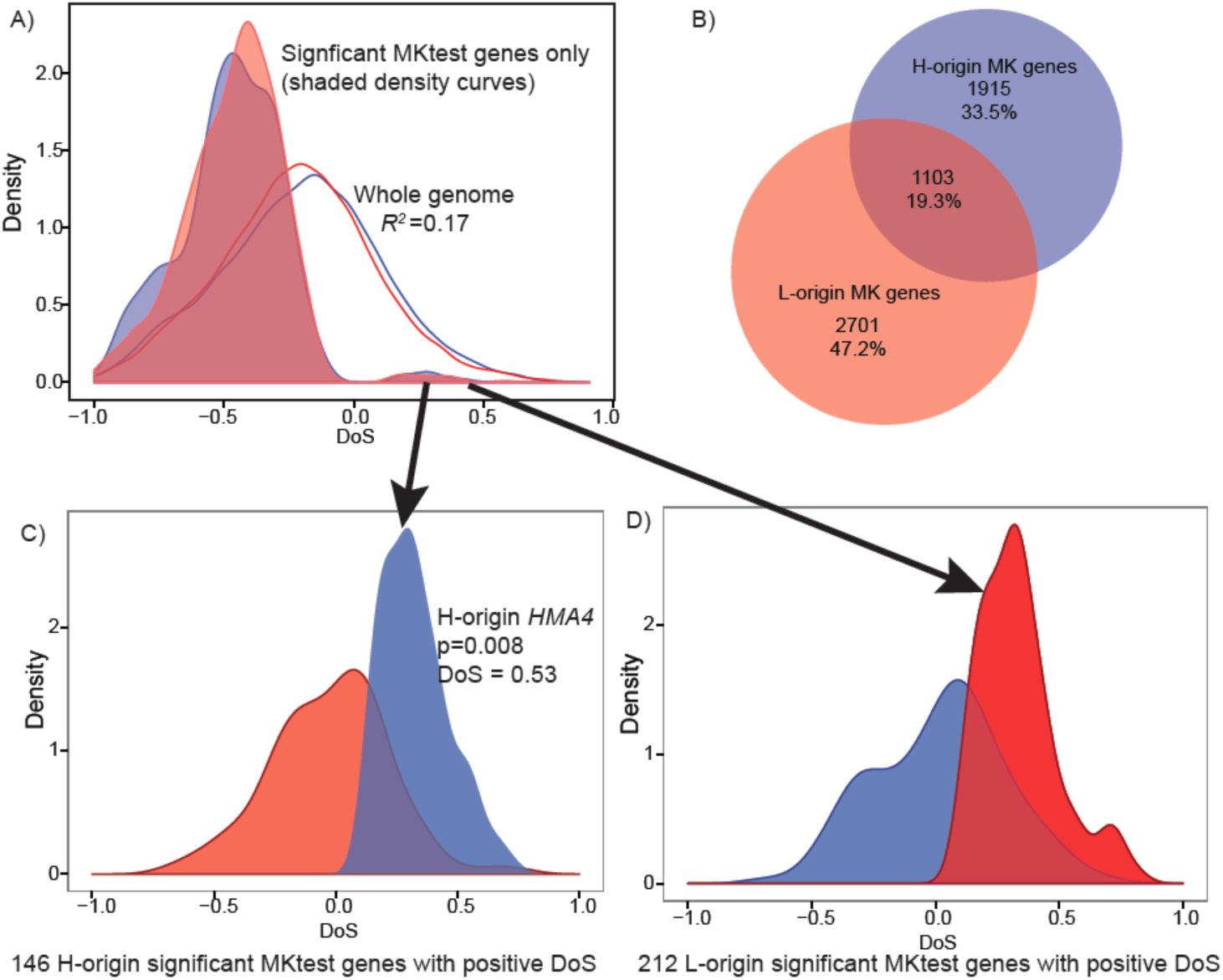
The direction of selection for both subgenomes. (A) Density curves of the direction ofselection (DoS)^14^ for about 21,000 coding sequences (blue line and density curves are DoS for H-origin genes, red line and density are DoS for L-origin genes). *Ne*utral genes are indicated by 0, while negative values indicate purifying selection and positive values indicate positive selection. The means of these distributions are -0.20 and -0.22 for the H− and L-origin homeologs respectively, and show that ~70% of both homeologs have a negative selection index (negative DoS). Shaded density curves are genes that were significant for MK-tests (p < 0.05 using Fisher’s marginal p-values). (B) Only 19% of genes show significance for MK-tests for both homeologs. (C) Using only significant MK-test genes with positive DoS for *halleri*-origin and (D) positive DoS for *lyrata*-origin genes show that the other homeolog has significantly more negative DoS (p-value < 2.2e-16 using pairwise t-test and Wilcoxon signed rank test) when one shows positive selection using comparisons of DoS distributions in both C and D.

MK-tests were conducted to detect homeologs showing purifying selection or adaptive evolution on amino-acid changing mutations. Among the significant MK-test genes, a total of 3018 H-origin and 3804 L-origin homeologs showed DoS < 0 (*D*_n_/*D*_s_ < *P*_n_/*P*_s_). This is consistent with purifying selection rather than positive selection for these genes. While the homeologs with significant MK-test comprise a substantial portion in our dataset, only 19% of them include both homeologs (i.e., there is significance for one homeolog but not the other for 81% of significant homeologous pairs, Fig. 3B). For example, the H-origin homeolog of the resistance gene *RPM1* (orthologous to *A. thaliana* gene: AT3G07040) was significant for the MK-test (DoS < 0) but the L-origin copy was not.

For genes showing positive selection (or adaptive evolution) using MK-tests, 146 *halleri*-origin and 212 *lyrata*-origin genes were significant when DoS > 0.01 (Fig. 3C, D). For these genes, when the *halleri*-derived homeologs shows a positive DoS, the *lyrata*-derived homeolog shows a more neutral or negative distribution in DoS and vice versa. Among these is the H-origin *HMA4* gene. These results, in addition to the low correlation in DoS between homeologous pairs and small overlap among all significant MK-test genes (Fig. 3B), indicates that a substantial proportion of homeologs have been shaped by different strengths of selection. These results are also in agreement with low correlations in Tajima’s D and Fay and Wu’s H despite for pairs of homeologs (Table 1), providing additional support that redundant genes exhibit significant differences due to stronger positive or purifying selection on only one of the two copies.

### The Distribution of Fitness Effects (DFE)

The tests above indicated that large numbers of homeologs show patterns consistent with purifying selection on amino-acid changing mutations (see Fig. 3). We quantified the genome-wide proportions of deleterious and effectively neutral mutations using the distribution of fitness effects (DFE) method^13^ in the two *A. kamchatica* subgenomes and both diploid relatives. In this method, the DFE is estimated from the site frequency spectra of non-synonymous and synonymous polymorphisms while accounting for effects of demographic changes. Effectively neutral mutations are represented by 0 < *N_e_s* < 1, mildly deleterious by 1< *N_e_s* <10, deleterious by 10 < *N_e_s* < 100 and strongly deleterious by *N_e_s* > 100 (where *N_e_* is the effective population size and *s* is the selection coefficient). The DFE estimates of the two *A. kamchatica* subgenomes show similar distributions with about 70% of mutations in the deleterious to strongly deleterious categories (*N_e_s* > 10) and about 20% effectively neutral (0 < *N_e_s* < 1) (Fig. 4A). The DFE of *A. halleri* and *A. lyrata* showed lower proportions of neutral mutations (16% of mutations 0 < *N_e_s* < 1 in diploids, and 19% mutations 0 < *N_e_s* < 1 in both subgenomes) and greater proportions of deleterious mutations (*N_e_s* > 100) than either of the corresponding allopolyploid subgenomes. While the differences are significant, the magnitude of the differences is not remarkable.

**Fig. 4.**
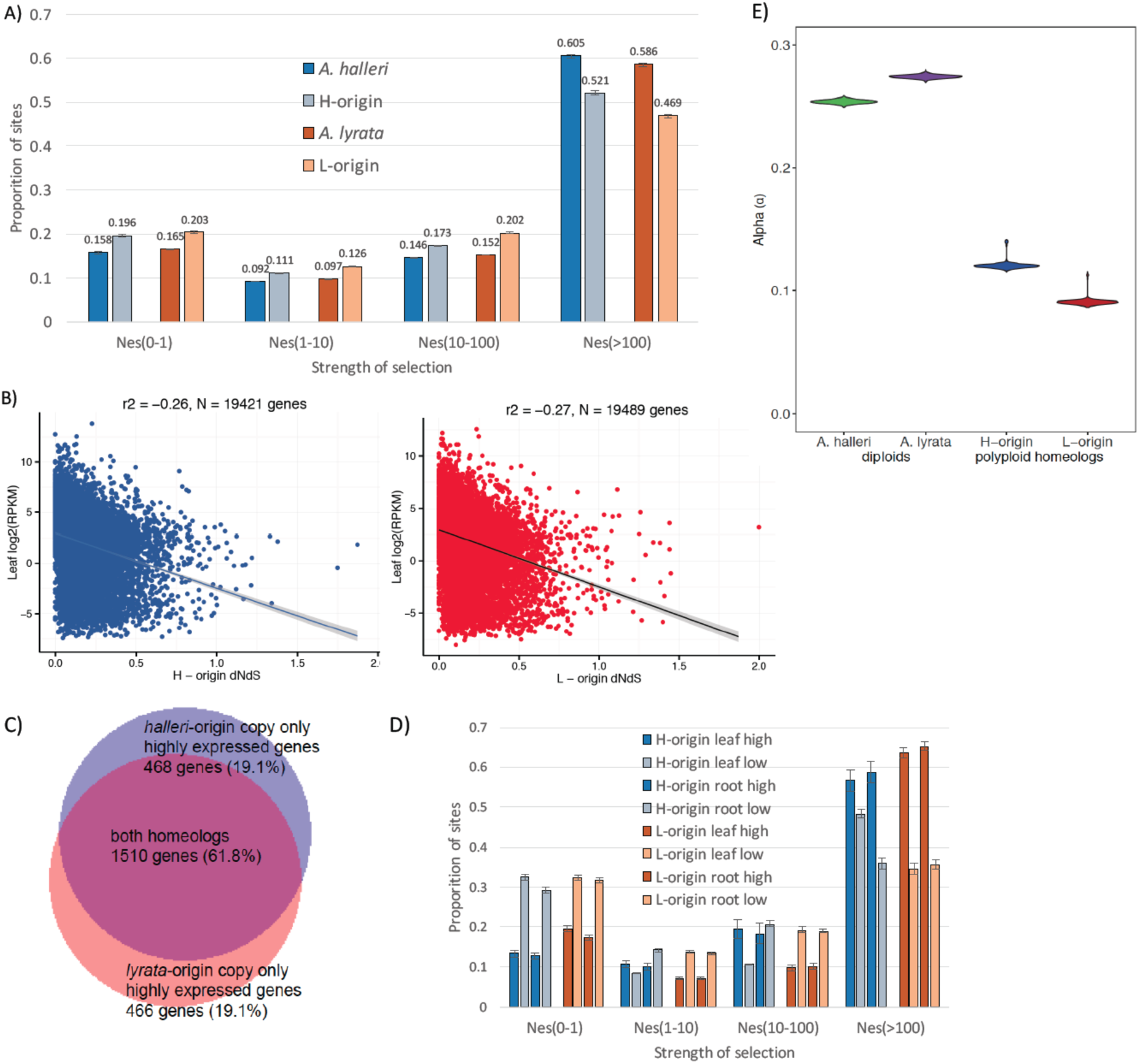
The strength of purifying selection and adaptive evolution. (A) The distribution of fitnesseffects (DFE) of deleterious mutations for coding sequences of the two *A. kamchatica* subgenomes and corresponding diploid orthologs of *A. halleri* and *A. lyrata*. The strength of selection is indicated by *N_
e_*s where *N_e_* is the effective population size and *s* is the selection coefficient. Error bars show standard deviations. (B) Evolutionary rates are negatively correlated with gene expression in both homeologs. (C) Overlap of genes that are highly expressed leaf tissues (in upper 10% of expression) in both homeologs. (D) DFE categorized by expression in both subgenomes. Expression categories were taken from the upper 10% (high) and lower 10% (low) of expression distribution in all *A. kamchatica* homeologs. (E) The proportion of adaptive substitutions (α) for both subgenomes (H-origin α = 0.12, CI: 0.117-0.141, L-origin α = 0.09, CI: 0.087-0.094) and for the two corresponding diploid species (*A. halleri* α = 0.25, CI: 0.251-0.257, *A. lyrata* α = 0.27, CI: 0.272-0.277) are significantly greater than zero.

To examine whether subsets of either subgenome experience a reduction in purifying selection, we classified homeologs according to gene expression level, which is one of the best predictors of evolutionary rates (dN/dS) in most organisms^49^. Expression level is negatively correlated with dN/dS due to strong constraint on amino acid substitutions (dN)^22^ for highly expressed genes, but this has not been shown in recent polyploid species. As a test of selective constraint on highly expressed genes, we found dN/dS was negatively correlated with expression for both homeologs (Fig. 4B). We would therefore expect genes that are highly expressed to show the strongest purifying selection, and low expressed genes to show relaxed constraint. We estimated the DFE again to quantify purifying selection and relaxed constraint using the distribution of expression levels in leaf and root tissues of *A. kamchatica* to categorize homeologs as high (genes in upper 10% RPKM) or low expression (lower 10% of RPKM). The majority (62%) of the highly expressed genes include both homeologs (Fig. 4C). The DFE patterns indicated that low expressed genes have the highest proportion of neutral mutations (relaxed constraint) and the lowest proportion of deleterious mutations compared with the genome-wide data, while highly expressed genes showed the opposite pattern (Fig. 4D). These results indicate that the DFE method can detect relaxed constraint and strong purifying selection as predicted when gene expression levels are accounted for.

### The Proportion of Adaptive Substitutions in Diploids and Allopolyploid Subgenomes

The proportion of adaptive substitutions (α) was estimated as the excess of between-species divergence relative to polymorphism as expected from the estimated DFE^13^ to account for slightly deleterious mutations. In contrast to the majority of the previously studied plant species including *A. thaliana*, we found significantly positive values of α for the two diploid species and both allopolyploid subgenomes. The diploid species *A. halleri* and *A. lyrata* showed the highest α values (0.25 and 0.27 respectively) (Fig. 4E). We subsampled 18 *A. kamchatica* accessions to be statistically comparable to the available *A. halleri* and *A. lyrata* samples (Supplementary Table 6). The α estimates for the H- and L-origin subgenomes of *A. kamchatica* were lower than those of the corresponding diploid species but significantly greater than zero (0.12 and 0.09, respectively) (Fig. 4E). The difference in α between subgenomes was significant but subtle (3% difference using the samples above, 6% difference when all 25 *A. kamchatica* accessions were used; Supplementary Fig. 4).

### High Impact Mutations are at Low Frequency in Subgenomes

We identified genes having high impact mutations that are likely to be deleterious due to their putative effects on amino acid sequences and gene expression into the following mutation categories: frameshifts, loss of start codon, premature stop codons (stop gained), and loss of stop codons (stop loss). For any gene, we counted every one of the mutation types regardless of the number. While it is not possible to determine the order of disruptive mutations, multiple frameshifts of premature stop codons in a gene would be expected to result in a loss of function.

Frameshifts and stop-gained categories comprised the majority of mutation types for both subgenomes (Supplementary Table 7). Frequencies of each mutation type indicated that most mutation types in any gene are found in only a single genotype in either subgenome (Fig. 5). Despite a higher number of mutations in the *lyrata*-homeologs, there were slightly greater proportions at low frequencies in the *halleri*-homeologs. Out the total 4219 *halleri*-origin and 4952 *lyrata*-origin disrupted genes, only 511 genes (2.5%) showed large effect mutations in both homeologs in the same accession suggesting that large effect mutations in both homeologs were deleterious. The distribution of genes with high impact mutations in both homeologs shows that most accessions have < 50 genes (orthologous to *A. thaliana*) that are disrupted with putatively similar functions (Supplementary Fig. 5).

**Fig. 5.**
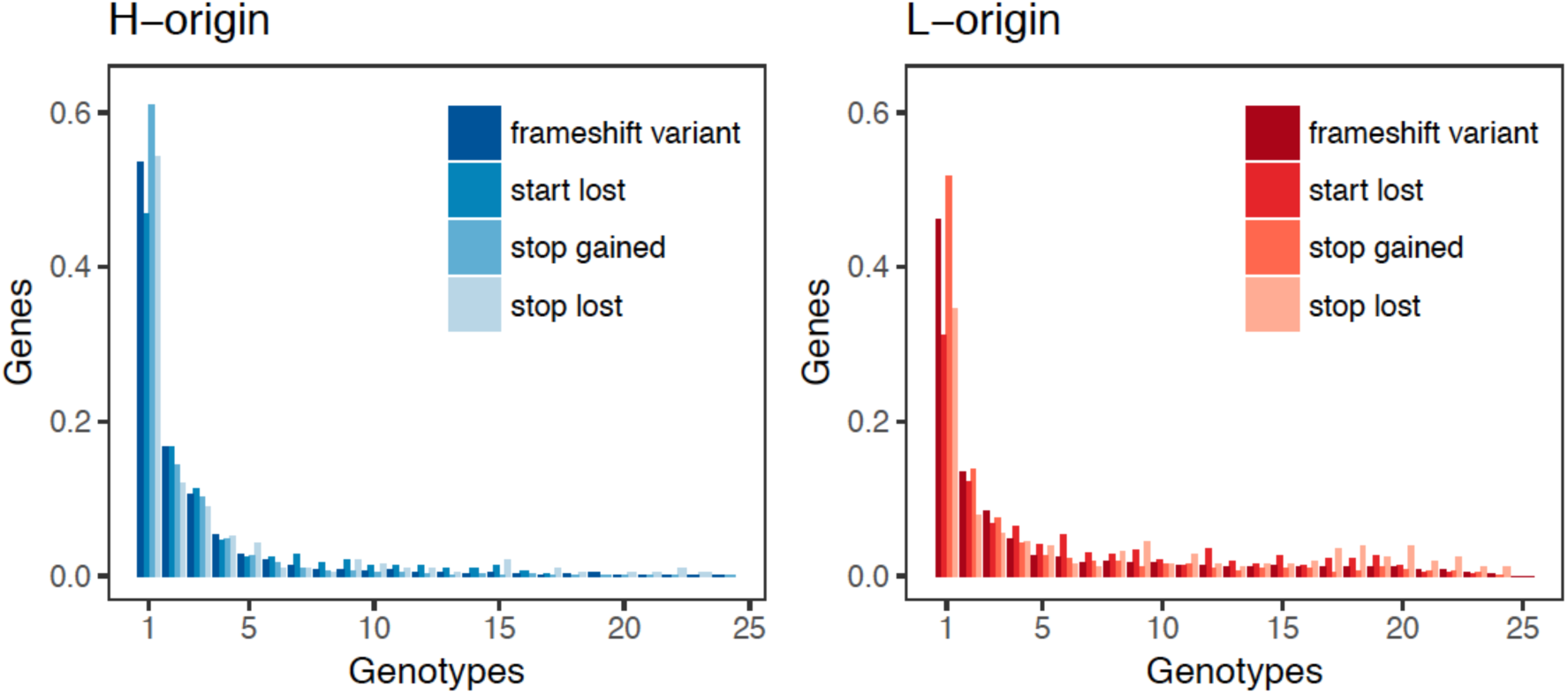
Frequency distributions of high impact mutations. Large effect mutations are at lowfrequency for both subgenomes.

We conducted gene ontology (GO) analysis to determine whether there was enrichment for GO terms using the two most common high-impact mutation types, i.e., frameshift mutations and stop codons. For both subgenomes, hydrolase activity (GO:0016787) was the most significant GO term for molecular function, followed by several GO categories for nucleotide binding (Supplementary Table 8). Programmed cell death (GO:0012501) and apoptosis (GO:0006915) were significant in the *halleri*-origin genes only. No significant gene ontologies were found with ≥ 20 query genes for the list of genes that had high impact mutations in both homeologs in a single accession.

## Discussion

### Similar Genome-wide Distributions in Both Subgenomes but Low Correlations Between Homeologous Pairs

A recurrent pattern we observed on patterns of diversity and signatures of selection was that the genome-wide distributions were similar between subgenomes, but the correlations between the pairs of homeologs were low. We found this pattern in the polymorphism levels such as π_syn_ and θ_w syn_, in frequency-based tests of neutrality (Tajima’s D, Fay & Wu’s H), and in divergence-based tests (DoS). The similar genome-wide distributions are consistent with the fact that the subgenomes shared the same history since the allopolyploidization event. The low correlation suggests that at the gene level, genetic diversity of a large number of homeologs may have been shaped by different levels of positive and purifying selection, as well as relaxed constraint. This supports that homeologs may evolve as independent loci, which may not be surprising because A. *kamchatica* shows disomic inheritance prohibiting recombination between homeologs^28^. These results also suggest that the difference between homeologs could contribute to the broad environmental response of polyploids, which may be realized by combining different adaptations of two parental species^10^ such as in the *HMA4* gene.

### Nucleotide Diversity and Linkage Disequilibrium is Similar to *A. thaliana* Suggesting the Feasibility of Genome-wide Association Studies of *A. kamchatica*

We found that the level of nucleotide diversity of *A. kamchatica* is moderate and similar to that of the diploid self-compatible *A. thaliana,* and 6 times lower than the diploid outcrossing species *A. halleri* and *A. lyrata*. It follows that the *N*_*e*_ of *A. kamchatica* is 6 times lower than the two diploid species. The ancestor of the genus *Arabidopsis* must have been a self-incompatible diploid species like present-day *A. lyrata* and *A. halleri*^25^, indicating that similar reductions in genetic diversity occurred in the lineages of *A. kamchatica* and *A. thaliana*.

The extent of LD decay in *A. kamchatica* is also comparable to *A. thaliana*^45^ and appears adequate for characterizing the genetic architecture of complex traits within relatively narrow genomic windows using genome-wide association studies (GWAS). The selfing mating system, levels of genetic diversity, LD, and a recently established transgenic technique^50^ suggests that *A. kamchatica* would be a suitable model for functional genomics of adaptive mutations in a polyploid species.

### The *HMA4* Locus Exhibits Significant Subgenome Differences in Genetic Diversity

The most important locus for zinc hyperaccumulation, *HMA4*^37^, involved two types of gene duplication in *A. kamchatica*: a tandem triplication in diploid *A. halleri*, followed by a whole genome duplication event, which contributed an additional *HMA4* copy from the *A. lyrata* parent. Despite multiple hybrid origins of *A. kamchatica*, the tandem triplication (three *halleri*-derived *HMA4* copies) is fixed in the allopolyploid^10^ suggesting it was present in all founding *A. halleri* parents. The high expression of H-origin *HMA4* in *A. kamchatica* explains high levels of zinc accumulation. Expression of the L-origin *HMA4* copy is very low compared with the *halleri HMA4* gene(s) so it is unlikely that the copy from *A. lyrata* contributes anything significant to hyperaccumulation in *A. kamchatica.*

Long scaffolds containing the *HMA4* copies and surrounding genes allowed us to compare homeologs across large genomic distances. The genetic diversity surrounding the *halleri*-derived *HMA4* gene that spans ca. 300 kb (HMA4-M) is significantly lower that the syntenic *lyrata*-derived region (ca. 100 kb) suggesting different evolutionary pressures or trajectories of functional duplicates. The higher diversity of the *lyrata* HMA4-M region is consistent with a pattern of relaxed constraint, while a selective sweep and genetic hitchhiking characterizes the *halleri-*derived HMA4-M region. Because we can infer that the triplication was ancestral and the reduced diversity at this locus and hitchhiking surrounding the *HMA4* genes was most likely the result of strong selection in the *A. halleri* parent^38^, diversity was probably greatly reduced prior to the polyploidization events.

### Purifying Selection in Polyploid Species

Theoretical studies suggested that higher proportions of neutral mutations (i.e., greater relaxed constraint) can result from whole genome duplication due to the reduction of *N_e_* or due to masking of deleterious mutations by functionally redundant gene copies^15,16^. This would be evident by greater proportions of effectively neutral mutations (0 < *N_e_*s < 1) in the polyploid subgenomes compared with the diploid parents^8^. Similarly, greater proportions of deleterious mutations (*N_e_*s > 10) in the diploid species would be expected compared to their derived polyploid subgenomes. We did detect significant differences between diploid parental species and the corresponding subgenomes of *A. kamchatica* in the proportions of mutations in the neutral (< 5% differences) and deleterious (5-7% differences) categories, although the differences were not drastic (Fig. 4A).

Using a similar approach, the change in purifying selection was studied by comparing the allopolyploid species *C. bursa-pastoris* with its diploid parents, *C. grandiflora* (outcrosser with high *N_e_*) and *C. orientalis* (selfing with low *N_e_*)^8^. First, for the subgenome derived from the outcrossing parent *C. grandiflora,* the proportion of neutral mutations doubled from ~17% to ~35% neutral mutations. This demonstrated that the subgenome derived from an outcrossing parent with a large *N_e_* shows a high proportion of neutral mutations due to relaxed constraint. Second, the opposite pattern was observed in the subgenome derived from the selfing parent (decreased from ~40% to 35% neutral mutations). The DFE patterns in *C. bursa-pastoris* and *C. orientalis* conforms to the trend in plants which shows species with low *N_e_* usually have greater proportions of neutral mutations^15^ consistent with greater strengths of purifying selection with higher *N_e_*. However, despite relatively high *N_e_* and outcrossing mating systems in *A. halleri* and *A. lyrata*, the differences in neutral mutations between the diploid species and the corresponding subgenomes are far less remarkable in *A. kamchatica* than in *C. bursa-pastoris*. These data suggest that *N_e_* alone is not adequate to explain the proportion of neutral mutations.

The strongest signal for relaxed constraint that we detected in the *A. kamchatica* subgenomes was observed when genes were categorized by expression levels. Genes that had low expression showed a significant increase in the proportion of neutral mutations (30-32%) over highly expressed genes (13-19%), and highly expressed genes show the strongest levels of purifying selection (for *N_e_*s > 10, 73-77% of mutations) in either subgenome. This result is consistent with expectations of stronger selective constraint on highly expressed genes^49^. A similar result was also found in the diploid *M. truncatula* where expression levels predicted very clearly the proportion of neutral mutations^51^, adding further support that the method is able to detect large differences in relaxed constraint when gene expression levels are taken into account.

Although theoretical analysis typically assumes that deleterious mutations may be masked by genome duplication, empirical studies showed that the dosage balance in gene networks may be a selective constraint^52^ and could work as a mechanism for purifying selection in an allopolyploid species. At this moment, the factors contributing to the difference between *A. kamchatica* and *C. bursa-pastoris* are not clear. It is possible that the time since the polyploidization events would not be adequate to detect the changes in the strength of purifying selection, although the time estimates of polyploidization overlap to a large extent (about 20,000-250,000 years ago for *A. kamchatica*, 100,000-300,000 years ago for *C. bursa-pastoris*).

### The Proportion of Adaptive Substitutions (α) are Significantly Greater Than Zero

This is the first report of α for *A. halleri* and *A. lyrata* using whole genome data, and to our knowledge, the first report of genome-wide α for a polyploid species. Previous multi-species comparisons showed that only a few plant species have α values that are greater than zero^17^, however these estimates were mostly done using limited genetic data (< 1000 loci)^17,19^ rather than genome-wide data. We estimated that 25-27% of non-synonymous substitutions are adaptive in the two diploid species *A. halleri* and *A. lyrata*. These are the highest estimates of α for any *Arabidopsis* species^17,40^ and higher than most plant species. The highest α among any plant species was estimated in the highly outcrossing *Capsella grandiflora* (α = 0.4-0.7)^19,53^ with levels similar to *Drosophila* and bacteria, all taxa with large effective population sizes^18^. Our results for the diploid species are consistent with previous studies that have shown a positive correlation between α and *N_e_* ^17,20^ which suggests that greater adaptive evolution often occurs in species with large effective population sizes, which is true for both highly outcrossing diploid species reported here.

Importantly, α for both subgenomes of *A. kamchatica* is also significantly greater than zero and indicates 6-12% of non-synonymous substitutions are adaptive. Many diploid plant species have a similar or larger effective population size than *A. kamchatica* (54,000-77,000), but did not show positive α^17^. For example, *N_e_* estimated for *A. thaliana* was between 65,000 – 267,000^17,20^ while α = -0.08^19^, indicating that effective population size alone cannot explain the significantly positive α of *A. kamchatica*. These data suggest that *A. kamchatica* has a positive α because of polyploidy. We suggest two mutually non-exclusive explanations. First, *A. kamchatica* may have inherited fixed non-synonymous or adaptive substitutions from the two parental species. The α values of *A. kamchatica* are roughly half of the parental species, in which the reduction may be attributable to the reduction of *N_e_*. Second, the rate of non-synonymous mutations are increased at the early stages of polyploid species in contrast to slow rate of old duplicated genes^21,22^. A classic idea of the high evolvability of duplicated genomes states that one of the duplicated copies may be able to obtain a new function or adaptive mutations because the other copy retains the original function^23,24^.

### High Impact Mutations with Deleterious Effects Were Rarely Fixed

The loss of homeologs in ancient polyploids, or nonfunctionalization, has been extensively studied ^24^, but relatively little is known about the population genetics of young polyploid species. We identified high impact mutations that are likely to disrupt the gene function. We found that about 20% of the homeologs in both subgenomes had disruptive mutations in our collection of 25 individuals (Supplementary Table 7), although their frequencies are low (Fig. 5) and only rarely are both homeologs disrupted. Interestingly, we found that high impact mutations were rarely fixed. This is in contrast with the results from another allopolyploid species *C. bursa-pastoris*, in which a large proportion of high-impact mutations (such as stop codon gained) were fixed^8^. In *A. kamchatica*, similar proportions of high-impact mutations were at low frequency compared with non-synonymous substitutions, which are also at low frequency (Fig. 1B), suggesting that genome-wide purifying selection keeps their frequency low, which is consistent with the prevalence of purifying selection shown by DoS and by DFE methods.

## Conclusion

Recently, new sequencing technology and algorithms drastically improved the genome assembly of crop polyploid species with a large genome size^54–56^ which will facilitate the genome-wide polymorphism analysis and scans for selection. By quantifying selection using polyploid species with different population sizes, times since polyploidization and mating systems, general patterns of selection in polyploid genomes will emerge. A further step will be to incorporate polymorphism, gene expression, and species distribution data (i.e., landscape genomics) of diploid parents and allopolyploid hybrids to identify the contributions of parental adaptations for broadening climatic regimes and abiotic habitats in polyploids.

## Materials and Methods

### Allopolyploid plant samples and resequencing

*Arabidopsis kamchatica* (Fisch. ex DC.) K. Shimizu & Kudoh^27^ is an allotetraploid species distributed in East Asia and North America. We consider Russian individuals described as *Cardaminopsis kamtschatika* or *Cardaminopsis lyrata* as synonyms (note that *Arabidopsis lyrata* is a distinct diploid species)^57^. Genomic DNA from 25 accessions of *A. kamchatica* was extracted from leaf tissue using the D*Ne*asy Plant Kit (Qiagen). These accessions were collected from Taiwan, lowland and highland regions of Japan, Eastern Russia, Sakhalin Island, and Alaska, USA (listed in Supplementary Table 3). DNA concentration and quality was measured using Qbit. Genomic DNA libraries were constructed at the Functional Genomics Center Zurich (FGCZ) using NEB *Ne*xt Ultra. Total DNA was sequenced on Illumina HiSeq 2000 using paired end sequences with an average insert size of 200-500 bp. Read lengths were 100 bp. For 22 accessions, a single lane included six *A. kamchatica* DNA samples and for three accessions (KWS, MUR, and PAK), eight samples per lane were used.

### Illumina read mapping and sorting using v2.2 reference genomes

Illumina reads from *A. kamchatica* were mapped using BWA-MEM version 0.7.10 on the two diploid genomes independently. We classified the reads to each parental origin as H-origin (*halleri*-origin) and L-origin (*lyrata*-origin) using HomeoRoq (http://seselab.org/homeoroq, last accessed July 14, 2016). In this method, reads from each accession were first mapped to each parental genome, and then classified as H-origin, L-origin, common, or unclassified (see fig. 1 in^32^ for schematic diagram). Here, the ‘common’ reads are the reads that aligned equally well to both parental genomes. After mapping to the *A. halleri* genome, we detected *A. kamchatica halleri*-origin (H-origin) reads and identified single-nucleotide polymorphisms (SNPs) and short insertions and deletions using GATK v3.3^58^. Then, the nucleotides were replaced on the detected variant position in the reference genome with the alternative nucleotides if the position (1) covered by at least 20% of the average coverage of reads in each library, (2) covered by at most twice of the average coverage and (3) has 30 or higher mutation detection quality (QUAL) produced by GATK. This cycle of mapping, read classification, and reference modification, was repeated ten times. For the reference modification, we used only *origin* reads the first five times and both origin and common reads the last five times. The *A. kamchatica lyrata*-origin (L-origin) genome was iteratively updated in a similar manner. The modified genomes were only used for read sorting. Coverage was calculated for both subgenomes of our resequenced lines by using the sum of the diploid parents as the genome size (250 + 225 = 475) and *common* plus sorted *origin* reads (Supplementary Table 3).

### Variant calling

For final variant calling, we combined the common reads of *A. kamchatica* with each sorted H-origin or L-origin reads and aligned them back to the original parental genomes using BWA-MEM v0.7.10. We called variants using GATK v3.3-0 following established best practices^59,60^. We processed each alignment BAM file separately to fix mate pairs, mark duplicates, and realign reads around indels. Then we identified variants by running HaplotypeCaller jointly on all genotypes but separately for each parental subgenome. To remove low-quality variants, we mostly used the thresholds recommended for variant data sets where quality score cannot be recalibrated ^60^. We applied quality by depth (QD < 2), mapping quality (MQ < 30), mapping quality rank sum (MQRankSum < -15) and genotype quality (GQ < 20) filters. Because some of our accessions had relatively low coverage, we considered that the recommended strand and read position filters might be too strict and we did not apply them. Finally, we removed all variants that GATK reported as heterozygous. We used diploid data from 9 accessions of European *A. halleri* and 9 accessions of European *A. lyrata* from Novikova et al.^30^ mapped to our diploid references genomes and called SNPs using the same criteria. The diploid VCF files were then phased using Beagle^61^ to produce 18 alleles for each species.

Regions with excessively high coverage are likely to be repetitive or incorrectly assembled, therefore variants called in those regions are probably spurious. To determine the coverage thresholds, we summed up the coverage reported by bamtools^62^ for each position in the final alignment files across all genotypes. We only considered reads with mapping quality (MQ) of at least 20. Then, we calculated the mean and standard deviation for the distribution of the obtained sums in each parental genome. We assumed a Poisson distribution and added 5 standard deviations to the mean to determine the thresholds. These thresholds (2891 and 2509 for *A. halleri* and *A. lyrata* respectively) were applied to the DP property (total depth of coverage across all genotypes) in the INFO field of the corresponding VCF file. In addition, we applied a coverage filter at genotype level to exclude calls with coverage below 2 or above 250.

To check for additional spurious variants, we randomly sampled 20 million reads (10 million per parent) from *A. halleri* and *A. lyrata* short-insert (200 bp) reads and ran it through the same variant calling pipeline as the *A. kamchatica* genotypes. The only difference between the runs was that this simulated sample was processed alone while variants for *A. kamchatica* genotypes were called jointly. Any variants called with the simulated sample would be due to incorrect read sorting between the parents or repetitive sequences present in the parental genomes. Such spurious variants would also be likely to appear among *A. kamchatica* variants even if the corresponding regions were completely conserved between *A. kamchatica* and its parents. Among the uncovered variants, 59,856 and 58,645 were also present in *A. kamchatica* on *A. halleri* and *A. lyrata* sides respectively. All of these variants were marked as filter failing. When applying polymorphisms to the reference sequences, we used N’s in positions where clear calls could not be made due to insufficient coverage, excessive coverage, low quality polymorphisms or heterozygosity. Such treatment allowed us to avoid using reference calls in regions where the actual sequence is highly uncertain.

### Coding sequence (CDS) alignments

We identified homeologous genes based on reciprocal blast hit (best-to-best with E-values < 10^−15^ and alignment length ≥ 200 bp) among coding sequences from the v2.2 *A. halleri* and *A. lyrata* genome annotations. Using the same approach, we also detected orthologous relationships between the predicted genes in diploid *A. halleri* and *A. lyrata* annotated genome assemblies and *A. thaliana* genes (TAIR 10). In cases of duplicated genes of interest such as *HMA4* (tandemly duplicated three times in *A. halleri*), we used only one copy for diversity analysis due to non-unique alignments of Illumina reads and very high sequence identity (99%) in the *A. halleri* reference genome. Therefore, our genome-wide dataset of coding sequences of homeologs do not contain genes that are duplicated in one genome but not the other.

To make coding sequence alignments, we individually applied SNPs and deletions from each of the 25 *A. kamchatica* genotypes (*H-origin* or *L-origin*) to the corresponding reference genomes. We omitted insertions in order to preserve the genomic coordinates of the coding sequences, which would consequently facilitate the alignment. If a variant was heterozygous, failed the genotype quality filter (GQ < 20), or was not called for a particular genotype (but called for other genotypes), the corresponding bases were replaced with N’s. We assumed that a sequence contains reference bases at positions that are not specified in VCF file and have adequate coverage. Therefore, all bases with coverage < 2 (insufficient) or > 250 (abnormally high) were replaced with N’s. After that, we extracted coding sequences from the modified genomes and grouped them by gene. Thus, each H-origin or L-origin gene had an alignment file containing 25 aligned coding sequences (one for each genotype). Finally, we aligned *A. thaliana* orthologs as an outgroup using Muscle v3.8^63^. With the profile alignment option, which preserved the alignment of the ingroup sequences and only aligned the outgroup sequence to the core ingroup alignment. The same procedure was used for making gene alignments of the 18 phased alleles for diploid *A. halleri* and *A. lyrata*.

### Population structure and phylogenetic analysis

We used 1000 randomly selected coding sequence (CDS) alignments from both *halleri* and *lyrata* derived homeologs. We then individually concatenated the *halleri* alignments and the *lyrata* alignments to use for population structure and phylogenetic analysis. The input data sets for the population structure analysis contained 21,341 and 16,223 markers from *halleri-* and *lyrata-*origin CDS respectively. We ran STRUCTURE v2.3.4^64^ ten times for each K = 1 to 9 using the admixture model and 50,000 MCMC rounds for burnin followed by 100,000 rounds to generate the data. The output was analyzed with STRUCTURE HARVESTER v0.6.94 and clusters were rearranged with CLUMPP v1.1.2. For phylogenetic analysis, we added *A. halleri* and *A. lyrata* as outgroups and ran Mr. Bayes v3.2.6^65^ with default parameters for 500,000 generations sampling every 1000^th^ generation.

### Coding sequence diversity and site frequency spectra

For gene alignments containing coding sequences, summary and diversity statistics, including divergence from *A. thaliana*, were estimated using *libsequence* packages ^66^ and custom R, Perl, and Ruby shell scripts. The *libsequence* programs *compute* and *Hcalc* were used to estimates average pairwise diversity (π), θ_w_, Tajima’s D, Fay and Wu’s H. Non-synonymous and synonymous diversity and gene based allele frequencies were estimated using the *polydNdS* program with the −P flag to generate SNP tables for each gene. The site frequency spectra (SFS), were created using the SFS.pl program available from the J. Ross-Ibarra (http://www.plantsciences.ucdavis.edu/faculty/ross-ibarra/code/files/ea3bd485e4c7dee37c59e8ba77ca800e-11.html) on the set of non-synonymous and synonymous polymorphisms identified using *polydnds*. Both folded and unfolded SFS were calculated; the folded spectrum does not differentiate between ancestral polymorphisms and polymorphism that are the result of mutations that have entered a population since it split from a common ancestor, while the unfolded spectra are based on derived allele frequencies. We converted the SFS data to SFS count tables using a custom python script (sfs_extraction.py). We used two published mutation rates, one based on the synonymous substitution rates calibrated by fossil records^41^, and another for total sites in mutation accumulation lines^42^, to estimate the effective population size using the following equation: *N_e_* = π_*syn*_ or π_total_ /4μ (where π was estimated from our data and μ from^41,42^).

### Linkage disequilibrium and sliding window diversity

To conduct sliding window analyses along entire scaffolds, we used the PopGenome R^67^ package to calculate diversity of all, intergenic, coding, exonic, and intron regions of *A. kamchatica* using *A. halleri* or *A. lyrata* derived VCF and reference gene annotation (.gff) files. We estimated the average nucleotide diversity, Watterson’s θ_w_ and π (the average number of pairwise nucleotide differences per site). To estimate genome-wide linkage disequilibrium (LD), we used the geno-r2 option in VCFtools^68^ across window sizes of a maximum distance of 20 kb, 50 kb or 1 Mb using a minor allele frequency ≥ 0.1, separately for the halleri or lyrata derived VCF files. The resulting r^2^ between SNPs were grouped into bins of 50 bp length. We estimated the average, 50% and 90% confidence intervals of correlation coefficients of each bin.

### Direction of selection (DoS), Distribution of Fitness Effects (DFE) and Adaptive Substitutions (α)

The program *MKtest* from the libsequence library, was used to count the total number of polymorphic non-synonymous (*P*_n_) and synonymous (*P*_s_) sites in *A. kamchatica* homeologs as well as the number of fixed non-synonymous (*D*_*n*_) and synonymous (*D*_s_) differences between *A. kamchatica* homeologs and *A. thaliana*. We used the program *MKtest* to perform standard tests on each gene for both homeologs separately; this is a contingency test comparing the numbers of between species difference and within species polymorphisms at non-synonymous and synonymous sites where significance is tested using Fisher’s exact tests for each gene.

Polymorphism and divergence data was used to calculate the direction of selection (DoS = *D*_n_/(*D*_n_+ *D*_s_) − *P*_n_/(*P*_n_ + *P*_s_)) statistic of ^14^. DoS < 0 is consistent with purifying selection and DoS > 0 is consistent with positive selection. To estimate the distribution of fitness effects (DFE, i.e. the distribution of the strength of selection acting against new mutations) and the proportion of adaptive substitutions (α) in *A. kamchatica*, *A. halleri* and *A. lyrata*, we used the likelihood method implemented in the software DoFE 3.0^13^. The program was run for 1×10^6^ steps, and sampled every 1,000 steps after a burn in of 100,000 steps. Strongly deleterious mutations have *N_e_^s^* > 10 (where *N_e_* is the effective population size and *s* is the selection coefficient), mildly deleterious mutations have 1 < *N_e_^s^* < 10, and effectively neutral mutations have *N_e_^s^* < 1. To estimate DFE we used folded allele frequency spectra and the estimated number of non-synonymous (*D*_n_) and synonymous (*D*_s_) differences between *A. kamchatica* homeologs or diploid orthologs and the corresonding outgroup *A. thaliana* orthologs.

### Transcriptome data

We used RNA-seq data collected from roots and leaf tissue of the *A. kamchatica* Murodo (Japan) and Potter (Alaska, USA) accessions from Paape et al. 2016 to calculate expression for all homeologs in our dataset. We mapped the RNA-seq data to v2.2 *A. halleri* and *A. lyrata* reference genomes and sorted the reads using method described in Akama et al. 2014. Thus, for each gene in our polymorphism dataset, we obtained expression data that is specific to either homeolog. We estimated expression levels using HTseq to count reads, then calculated reads per kilobase of transcript per million mapped reads (RPKM). The mean RPKM values from three libraries of both leaf and root were used to make a distribution in RPKM that corresponds to our polymorphism gene dataset. The distribution of RPKM was used to determine the upper and lower 10% tails in expression for both homeologs separately.

### Detection of High Impact Mutations

We used SnpEff v4.2^69^ to detect genetic variants that have putative loss of function mutations in both subgenomes of *A. kamchatica*. We ran the program separately on the variant file of each subgenome. First, we built custom databases for each parental genome using our v2.2 parental assemblies and annotation. Since SnpEff ignores filter fields in VCF files, we have removed all variants that failed our filters, replaced all genotypes that failed genotype filters with no-calls (i.e. ‘./.’), and removed any entries without valid variant calls. Such filtering allowed us to extract accurate gene summaries from SnpEff output.

SnpEff annotated polymorphisms within 32,410 and 31,119 genic regions in *A. halleri* derived and *A. lyrata* derived genomes respectively. These include all mutations with any impact type, but we focused only on frameshifts, premature stop codon, loss of stop codons and loss of start codons. The gene sets were thus reduced to 31,193 and 31,119 genes for *A. halleri* and *A. lyrata* derived genomes respectively. There are 21,419 and 21,463 reciprocal best BLAST hits between respectively *A. halleri* or *A. lyrata* and *A. thaliana*. Based on the intersection of these two data sets, we identified 20,292 homeologs between *A. halleri* and *A. lyrata*. Out of these 19 and 18 *halleri*-origin and *lyrata-*origin genes had no coverage. Gene ontology (GO) analysis was perfomed using agriGO (bioinfo.cau.edu.cn/agriGO) conducted using a custom annotation containing 19,936 GO annotations that correspond to *A. thaliana* orthologs with reciprocal-best BLAST hits for both homeologs. We used only queries with at least 20 genes.

## Acknowledgments

We thank Takashi Tsuchimatsu, Polina Novikova and Peter Keightley for useful discussions, and the Functional Genomic Center Zurich for sequencing services and technical support. The study was supported by Swiss National Science Foundation, the University Research Priority Program of Evolution in Action of the University of Zurich, JST CREST Grant (number JPMJCR16O3) to KKS, MEXT KAKENHI Grant Number 16H06469, 26113709, Young Investigator Award of Human Frontier Science Program to KKS and JS, European Union’s Seventh Framework Programme for research, technological development and demonstration under grant agreement no GA-2010-267243 – PLANT FELLOWS to RVB and TP, Marie-Heim Hoegtlin grant by Swiss National Science Foundation to RSI, ISCB (Indo-Swiss Collaboration in Biotechnology) to KKS and MH, the Special Coordination Funds for Promoting Science and Technology from MEXT Japan, an Inamori Foundation research grant, a Japan Society for the Promotion of Science Grant-in-Aid for Scientific Research (Young Researchers B, 2277023), and Research and Education Funding for Japanese Alps Inter-Universities Cooperative Project, MEXT, Japan to KT.

## Data Accessibility

Illumina reads submitted to DDBJ. BioProject Submission ID: PSUB006170

The Sanger sequences were submitted to GenBank. GenBank BankIt submission. Submission ID: 2025864

Code for A. lyrata genome assembly https://gitlab.com/rbrisk/AlyrAssembly

Code for Variant calling in *A. kamchatica* https://gitlab.com/rbrisk/AkamVariants

